# Spatial updating of attention across eye movements: A neuro-computational approach

**DOI:** 10.1101/440727

**Authors:** Julia Bergelt, Fred H. Hamker

**Affiliations:** Artificial Intelligence, Department of Computer Science, Chemnitz University of Technology, Chemnitz, Germany

**Keywords:** eye-movement, neuro-computational model, space perception, predictive remapping, spatial updating of attention

## Abstract

While scanning our environment, the retinal image changes with every saccade. Nevertheless, the visual system anticipates where an attended target will be next and attention is updated to the new location. Recently, two different types of perisaccadic attentional updates were discovered: Predictive remapping of attention before saccade onset (Rolfs, Jonikaitis, Deubel, & Cavanagh, 2011) as well as lingering of attention after saccade (Golomb, Chun, & Mazer, 2008; Golomb, Pulido, Albrecht, Chun, & Mazer, 2010). We here propose a neurocomputational model located in LIP based on a previous model of perisaccadic space perception (Ziesche & Hamker, 2011, 2014). Our model can account for both types of updating of attention at a neural systems level. The lingering effect originates from the late updating of the proprioceptive eye position signal and the remapping from the early corollary discharge signal. We put these results in relationship to predictive remapping of receptive fields and show that both phenomena arise from the same simple, recurrent neural circuit. Thus, together with the previously published results, the model provides a comprehensive framework to discuss multiple experimental observations that occur around saccades.

## 1 Introduction

During natural vision, scene perception depends on exploratory scanning using accurate targeting of attention, saccadic eye movements, anticipation of the physical consequences of motor actions, and the ability to continuously integrate visual inputs with stored representations of previously viewed portions of the scene. For example, when there is an impending eye movement, the visual system can anticipate where the target will appear on the retina after the eye movement and, in preparation for this, spatial attention updates and moves to a new location. When subjects are instructed to monitor a particular location in the scene while moving the eyes, two different types of spatial attention shifts were recently discovered. One study shows that after a saccade, spatial attention lingers at the (irrelevant) retinotopic position, that is, the focus of attention appears to shift with the eyes but updates to its original world-centered position only after the eyes land at the saccade target location (Golomb et al., 2008, 2010). Another study by Rolfs et al. (2011) shows, that shortly before saccade onset, a locus of attention appears at a position opposite to the direction of the saccade, which suggests an anticipatory correction of the effects of eye movements. While these results initially appear contradictory, Jonikaitis, Szinte, Rolfs, and Cavanagh (2013) show, that both updating mechanisms occur simultaneously. Around the time of an eye movement, they detected attentional effects at two different locations at the same time although only one location was cued. This suggests, that there must be at least two attention pointers in addition to the saccade target location that are active around saccades. While the anticipatory shift of attention opposite to saccade direction may be potentially useful, although a bit too early, the shift of attention in the direction of the saccade may appear an error or at least a delay in the spatial updating. It is not clear, if the recently observed phenomena of spatial updating of attention relate to the observation of predictive remapping of receptive fields. In the seminal study of Duhamel, Colby, and Goldberg (1992), a flashed stimulus in the future receptive field, i.e. the location of a neuron’s receptive field after saccade, evoked a neural response prior to saccade. While this can be interpreted as an anticipatory shift of the receptive field, Cavanagh, Hunt, Afraz, and Rolfs (2010) suggested that it may also be explained by learned horizontal or lateral connections. Furthermore, they proposed that such a transfer of activation may also be responsible for spatial updating of attention.

Recently, Ziesche and Hamker (2011) proposed a model of perisaccadic perception to describe the underlying mechanisms of perceiving a stable world during eye movements. It uses gain fields and radial basis functions to perform coordinate transformations between different frames of references, more precisely between eye-and head-centered reference frames, which may possibly take place in the parietal cortex, specifically the lateral intraparietal cortex (LIP). The model accounts for predictive remapping using two eye position related signals, a discrete eye position signal and a corollary discharge signal, to compute the perceived position of stimuli across saccades. Ziesche and colleagues demonstrated the model is in addition able to explain the perisaccadic mislocalization of briefly flashed stimuli in complete darkness (Ziesche & Hamker, 2011) and the observation of saccadic suppression of displacement (SSD) (Ziesche & Hamker, 2014; Ziesche, Bergelt, Deubel, & Hamker, 2017).

We here will explore by means of the neuro-computational model the relationship between predictive remapping of receptive fields (Duhamel et al., 1992) and predictive remapping of attention. How do these phenomena occur at the neural systems level? May both recruit the same neural mechanisms? If yes, why does the attention pointer update opposite to saccade direction, while the receptive fields update with the saccade vector? Furthermore, we address the question if SSD and perisaccadic mislocalisation in complete darkness rely on similar mechanisms as the updating of attention pointers.

## 2 Neuro-computational Model

Ziesche and Hamker’s model simulated actual experiments using a simplified one-dimensional design. To allow for realistic simulations of more complex experiments we started by extending the original model to work in two dimensions. This two-dimensional model uses the same input signals, interactions and concepts as the one-dimensional model proposed by Ziesche and Hamker (Ziesche & Hamker, 2011, 2014). In addition to the existing inputs, we expanded the model by including a fourth input signal to introduce top-down attention to the system. We will now summarize briefly how the principles of the model work together and point out the parts where the translation from 1D to 2D and the extension take effect.

Like the original model, our new 2D model is based on two extraretinal, eye position related oculomotor signals, namely the proprioceptive eye position (PC) signal and the corollary discharge (CD) signal. The former signals current eye position and the latter saccade related motor components. The model is inspired by gain fields and radial basis function networks to perform coordinate transformations between eye-centered and head-centered reference frames (Pouget, Deneve, & Duhamel, 2002). However, unlike previous models, our 1D and 2D model account for the full temporal dynamics around saccades. Figure 1 shows the structure of the new 2D model with its maps and their way of interaction. The proprioceptive eye position signal (PC signal) encodes the eye position in head-centered coordinates and is proposed to originate in the primary somatosensory cortex (SI) (Wang, Zhang, Cohen, & Goldberg, 2007) or in the central thalamus (Tanaka, 2007) and is represented as *Xe_PC_*. In monkeys, the eye position signal has been shown to update around 60ms after saccade completion (Y. Xu, Wang, Peck, & Goldberg, 2011). In humans, updating latencies have not been measured, but there is evidence that updating latencies may be species specific, depending on the behavioral task. For example, in the SSD task data of monkey and human appears to differ (Joiner, Cavanaugh, FitzGibbon, & Wurtz, 2013). The corollary discharge (CD), which encodes the eye displacement retinotopically as it is a copy of the oculomotor command, is only active around saccade onset. It is likely routed from the superior colliculus (SC) via the mediodorsal thalamus (MD) to the frontal eye field (FEF) (Sommer & Wurtz, 2004), where it is transferred into a head-centered reference frame (Cassanello & Ferrera, 2007) representing the expected postsaccadic eye position. Both eye related signals are encoded in a two-dimensional spatial map representing horizontal and vertical dimension of a visual scene. The PC signal is used to transfer the eye-centered CD signal encoded in the eye-centered map *Xe_CD_* into a head-centered signal. Following the concept of coordinate transformation as suggested by Pouget et al. (2002), the gain fields that multiplicatively combine the CD and PC signal are required to be represented – from a computational point of view – in a four-dimensional representation, as computed in *Xe_FEF_*. The stimulus position is encoded in a two-dimensional retinocentric reference frame *Xr* modeling early extrastriate visual areas like MT or V4. The retinal signal feeds into two maps assumed to be located in the lateral intraparietal area (LIP) where it is gain modulated by either the eye position signal or the corollary discharge signal to obtain a joint representation of the stimulus position and the eye position or eye displacement, respectively. As the two LIP maps, called *Xb_PC_* and *Xb_CD_* according to the modulating signal, combine two two-dimensional signals multiplicatively, they again – from a computational point of view – have to be designed as four-dimensional maps. Both LIP maps interact with each other via intermediate neurons organized in a two-dimensional head-centered reference frame *Xh* using feedback projections. These intermediate neurons combine the information from both LIP maps and as a result encode the perceived spatial position of a stimulus in a head-centered reference frame.

**Figure 1:**
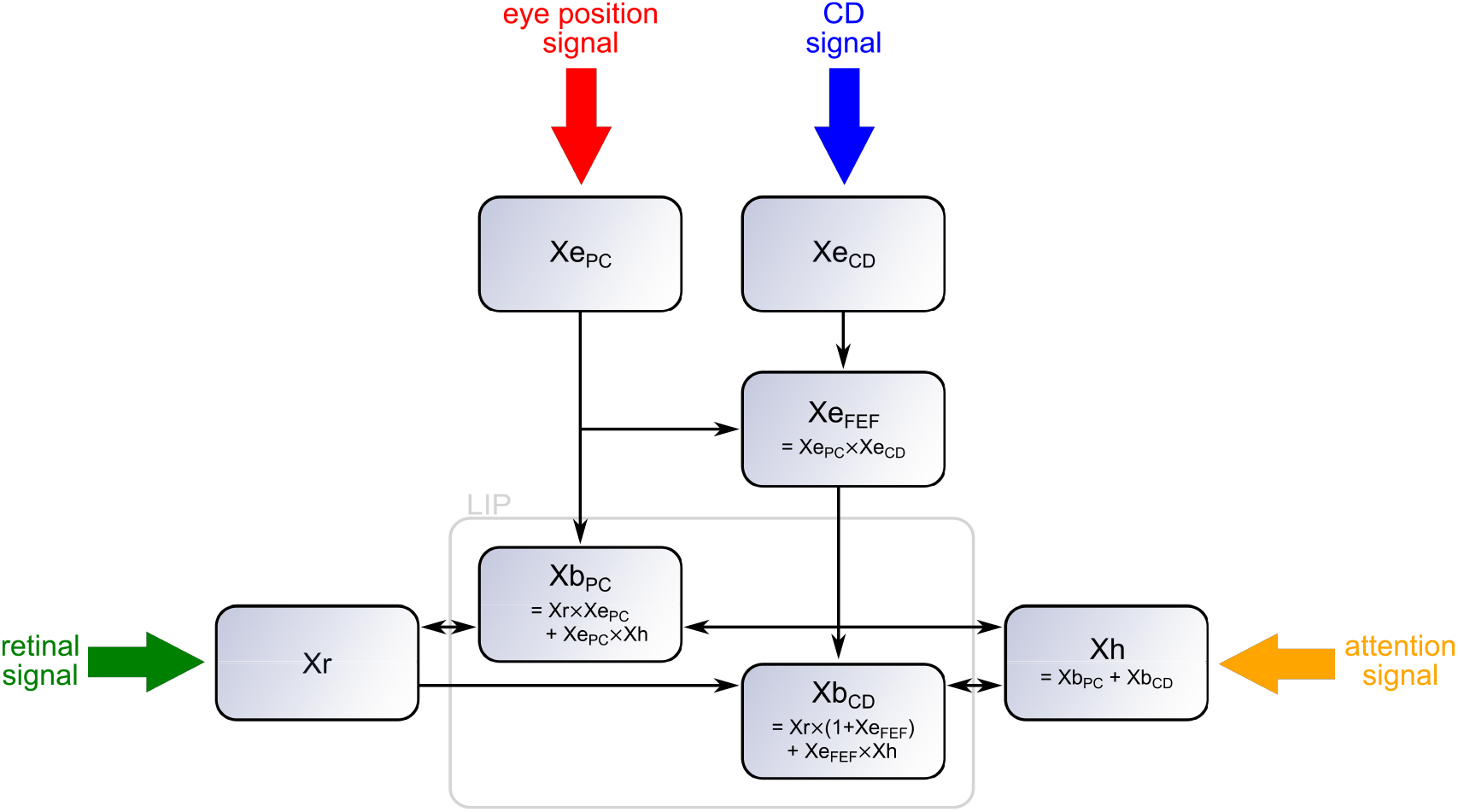
Structure of the neuro-computational model. The four input signals (retinotopic stimulus position *Xr*, proprioceptive eye position (PC) *Xe_PC_*, corollary discharge (CD) *Xe_CD_* and attention *Xh*) are fed into two LIP maps (*Xb_PC_*, *Xb_CD_*) which are gain modulated by either the PC signal or the CD signal. While the PC signal encodes the eye position in head-centered coordinates, the CD signal is originally eye-centered and must be first transferred into a head-centered reference frame. This is done in map *Xe_FEF_* using the eye position signal from *Xe_PC_*. The activities of all simulated LIP neurons are combined in map *Xh*, and from there feed back into both LIP maps. The interaction of the maps are summarized in the structural equations. The mathematical description of the model including all equations can be found in the Appendix.

Following the idea of LIP as a priority map (Goldberg, Bisley, Powell, & Gottlieb, 2006; Bisley, 2011), any activation in LIP is interpreted as an attentional signal and we refer to this as an attention pointer. In order to simulate the dynamics of spatial attention around eye movements, we can apply the model in two operational modes: a bottom-up mode where we cue attention by a stimulus and observe how the initially cued activity evolves during saccade, and a top-down mode where we set a head-centered top-down pointer into the model by providing a static signal to *Xh*. Given this static attention signal, we will read out the dynamics of attention while being transformed by the two eye position signals traveling through the feedback pathways towards an eye-centered representation of spatial attention.

## 3 Results

### Predictive remapping

Predictive remapping is a phenomena where the retinotopic position of a spatial restricted neuronal receptive field (RF) shifts in anticipation of a saccade such that the neuron responses to stimuli in the future receptive field (FRF) before the eye movement onset. Predictive remapping was first observed in LIP by Duhamel et al. (1992) and has been proposed to play a key role in establishing the percept of a visually stable world (Sommer & Wurtz, 2006; Wurtz, 2008; Mathôt & Theeuwes, 2010). Predictive remapping has been observed in many other areas: in extrastriate visual cortex (Nakamura & Colby, 2002), SC (Walker, Fitzgibbon, & Goldberg, 1995) and FEF (Umeno & Goldberg, 1997; Sommer & Wurtz, 2006). However, in these early studies predictive remapping was not clearly delineated from other receptive field shifts as discussed by Zirnsak, Lappe, and Hamker (2010) which triggered new experimental observations (Zirnsak, Steinmetz, Noudoost, Xu, & Moore, 2014; Neupane, Guitton, & Pack, 2016a, 2016b; Hartmann, Zirnsak, Marquis, Hamker, & Moore, 2017).

This report focuses on modeling the neural substrates of the original predictive remapping phenomenon, as described by Duhamel et al. (1992). In our model, predictive remapping arises from the feedback connection from *Xh* to *Xb_CD_*. To demonstrate this, we conducted two simple experiments, similar to those of Duhamel et al. (1992). In the first experiment, called fixation task, a stimulus is presented in the receptive field (RF) for 100ms while the eyes remain fixed at a particular location (FP at (0*°,* 0°)). The second experiment is called saccade task, as a saccade is executed from the fixation point (FP) to the saccade target (ST) at (10°, 8°). 150ms before saccade onset, a stimulus is presented for 100ms in the future receptive field (FRF), which is at the position of the receptive field after the eyes landed at the saccade target. The eye movement is simulated with the model of Van Wetter and Van Opstal (2008) and for this saccade length, it was 65ms in duration. The setup as well as the positions of the three input signals eye position, corollary discharge and retinal signal are shown in Figure 2.

**Figure 2:**
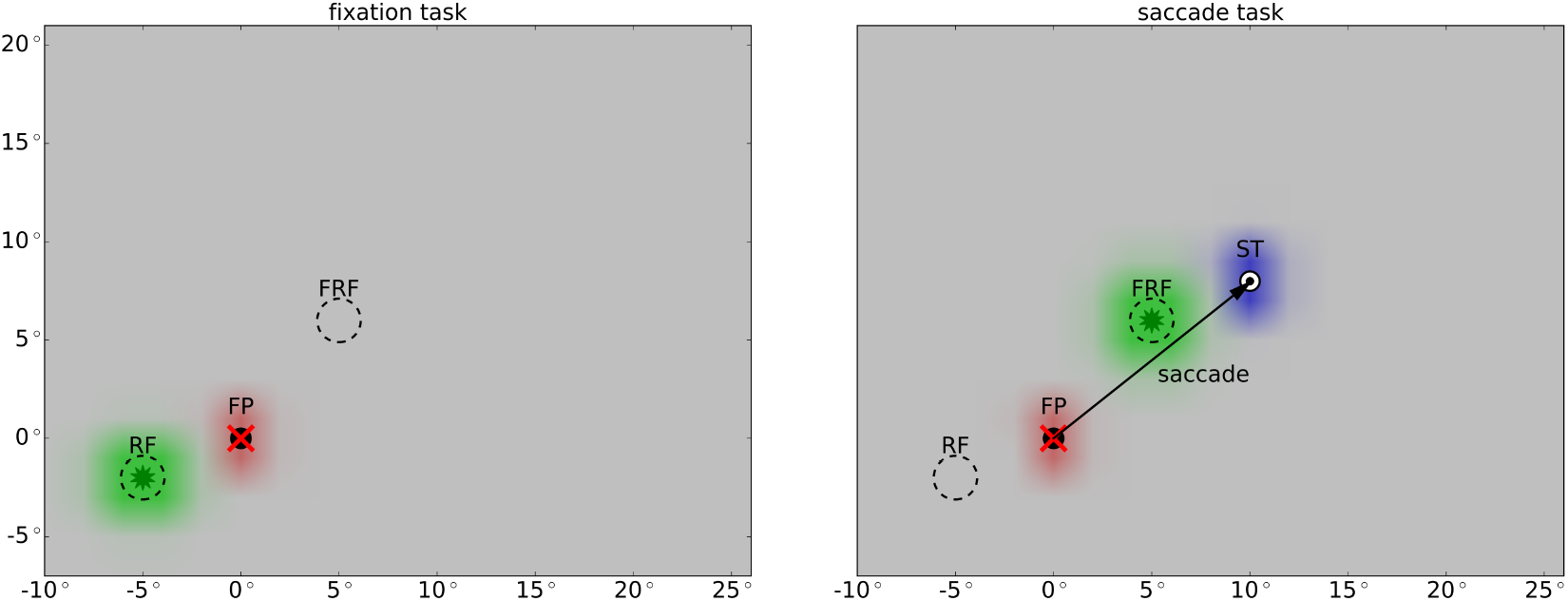
Setup for predictive remapping with respect to the three input signals (cf. Figure 1): In the fixation task (left), the eyes fixate at the fixation point (FP) at (0°, 0°) for the whole time. A stimulus (depicted by a green star) is presented in the receptive field (RF) at (−5°, −2°) for 100ms. In the saccade task (right), the stimulus is presented in the future receptive field (FRF) at (5°, 6°) 150ms before saccade onset for 100ms. The eye movement from fixation point (FP) to saccade target (ST) at (10°, 8°) lasts 65ms. The red cross symbolizes the current eye position. The colored blobs represent neural activity of the three different signals which change with time. The retinal signal (green) is delayed for 50ms to mimic the response latency. The eye position signal (red) starts to update 32ms after saccade offset to the new location of the eyes. The CD signal (blue) rises 86ms prior to saccade onset and is active for 171ms with its peak activity reached 10ms after saccade onset. As the retinal signal and the origin of the corollary discharge signal are retinotopic, they shift with the eye movement. In contrast, the PC signal is head-centered and therefore fixed during the saccade.

The results of the simulations are shown in Figure 3. To visualize the 4D LIP maps, we projected each map to two two-dimensional planes representing either the horizontal or the vertical information. Further, we show the neural activities of both LIP maps projected into the retinotopic space together with the spatial setup. In the fixation task (Figure 3A), a stimulus is presented in the receptive field, no saccade is planned and the eyes fixate at the fixation point (FP). The LIP map *Xb_PC_* combines the eye position signal at (0°, 0°) and the retinal signal at (*−*5°*, −*2°) multiplicatively, thus resulting in a single locus of activity at the crossing of both signals (at (0°*, −*5°) for horizontal and (0°*, −*2°) for vertical, respectively). As the retinal signal feeds continuously into the LIP map *Xb_CD_*, we get an activity line along one axis at the position of the stimulus (at 5° for horizontal and 2° for vertical, respectively). In the saccade task (see Figure 3B), the CD signal rises shortly before saccade onset and modulates the retinal signal of the stimulus presented in the future receptive field in such a way that it increases the neural gain at a single blob where the CD signal position crosses the retinal signal (at (10°, 5°) for horizontal and (8°, 6°) for vertical, respectively). The activity in LIP PC is the result of the multiplicative interaction of the PC signal and the retinal signal (at (0°, 5°) for horizontal and (0°, 6°) for vertical, respectively).

Activity along each diagonal of *Xb_PC_*, e.g. along the yellow dashed line, is read out and fed to one neuron in *Xh*. Likewise, the activity in the LIP CD map is read out along each diagonal and projected to *Xh*. At the same time, the activity of *Xh* is fed back to both LIP maps along each diagonal, which allows an interaction between *Xb_PC_* and *Xb_CD_* via *Xh*. In LIP CD, the feedback from *Xh* is combined multiplicatively with the CD signal. This leads to a second locus of activity in LIP CD (at (10°, −5°) for horizontal and (8°, −2°) for vertical, respectively). Note, that the position of this second locus of activity has the same “visual” position as if we presented the stimulus in the receptive field (cf. Figure 3A). That means, the neurons in *Xb_CD_* anticipate the updating of the stimulus position from FRF to RF with the help of the CD signal and the feedback from *Xh*. Thus, the *Xb_CD_* neurons show predictive remapping. In contrast, the LIP map for the PC signal has no such predictive component and contains only one single activation blob that represents the stimulus position with respect to the current eye position.

**Figure 3:**
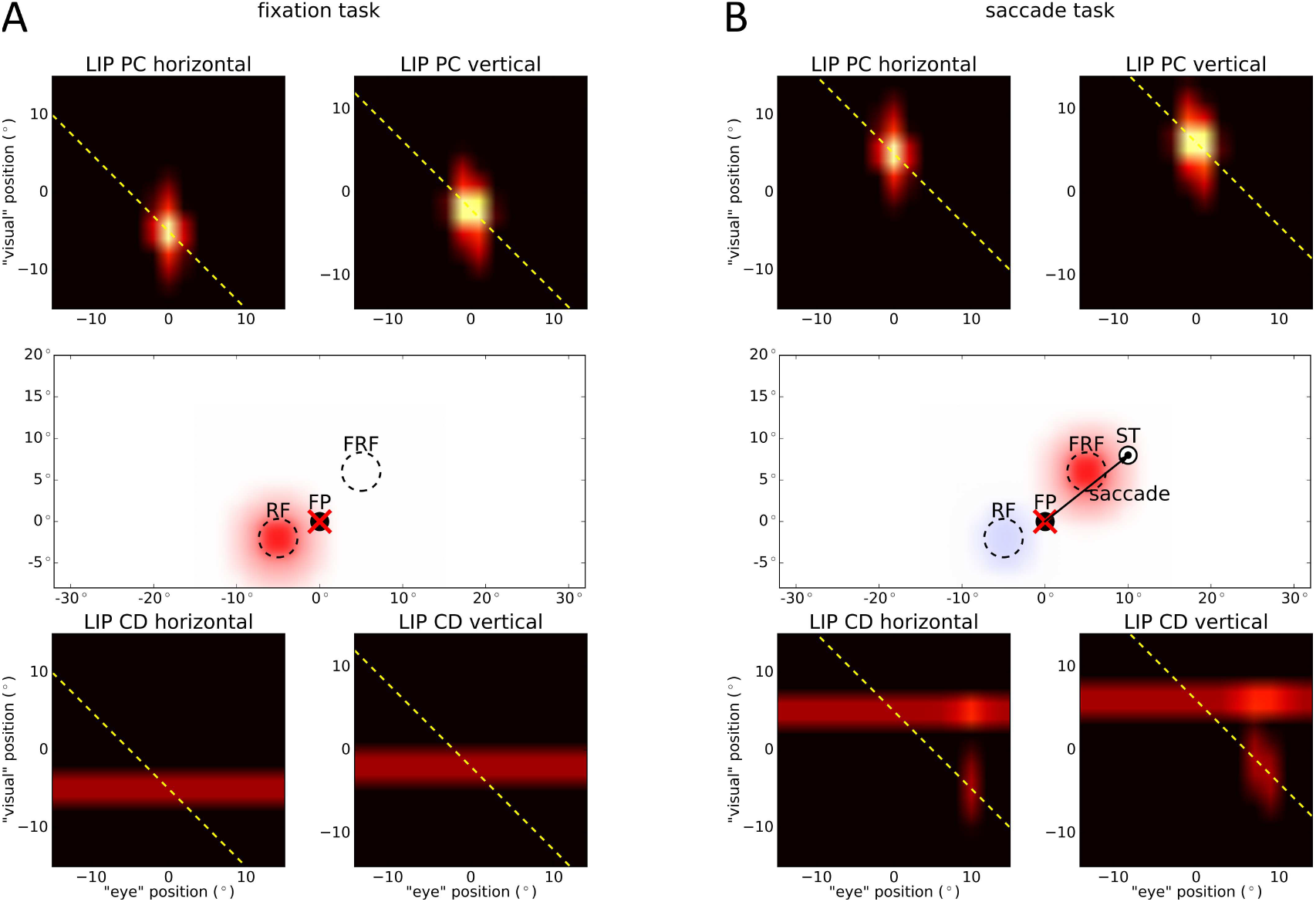
Simulation results of the two predictive remapping tasks. The activity of both LIP maps projected onto two two-dimensional planes are plotted representing horizontal and vertical information. The symbols are identical to Figure 2. (A) In the fixation task, a stimulus is presented in the receptive field (RF), the eyes fixate at FP and no saccade is planned. Due to the multiplicative interaction of eye position signal and retinal signal representing the stimulus, there is a single activation blob in LIP PC. Projected to visual space, this results in activity at RF (red blob). In LIP CD, there is an activity line indicating the position of the stimulus, which leads to activity at RF, too (covered by red blob). (B) In the saccade task, a stimulus is presented in the future receptive field (FRF) and a saccade is going to be executed from FP to ST. In LIP PC, there is an activity blob at the crossing of the current eye position and the stimulus position similar to the fixation task. Thus, we see activity at FRF triggered by LIP PC (red blob). Shortly before saccade onset, the CD signal rises which produces an additional peak of activity along the activity line from the stimulus signal at the position of the saccade target (ST) in LIP CD. Additionally, there is a second activity blob resulting from the interaction of the CD signal with the feedback signal from *Xh* along the yellow dashed diagonal. This leads to a second activity blob at RF (blue blob).

To summarize, our model suggests that predictive remapping in LIP is generated with the help of the corollary discharge signal. When a stimulus is presented, the eye-centered retinal signal feeds into both LIP maps. Thus, neurons whose receptive fields match with the stimulus position become active. Long before a saccade is planned the eye-centered stimulus position (from *Xr*) is transformed into a head-centered position through *Xb_PC_* with the help of the eye position signal and is encoded in *Xh*. Shortly before saccade onset, the CD signal rises and with the feedback from *Xh*, it activates neurons in *Xb_CD_* which encode the stimulus with respect to the future eye position. Thus, if the stimulus presented in the FRF is encoded with reference to ST, this stimulus – in a retinocetric reference frame – leads to a neural activation at RF as the eyes are still at FP and consequently, predictive remapping emerges. Predictive remapping does not require all-to-all connections, only a diagonal connectivity scheme within a small recurrently connected network of neurons.

### Spatial updating of attention

We particularly compare our simulation results to the data of Jonikaitis et al. (2013) who demonstrated both types of attentional updating in their data: Predictive remapping of attention reported by Rolfs et al. (2011) and lingering of attention reported by Golomb et al. (2008, 2010). Predictive remapping of attention occurs shortly before saccade onset and spatial attention is remapped to a position opposite to the direction of the saccade. In contrast, Golomb et al. reported that after a saccade, spatial attention lingers at the (irrelevant) retinotopic position, that is, the focus of attention shifts with the eyes but updates to the original fixed world-centered position only after the eyes landed. Using our model we replicate the observation of Jonikaitis et al. (2013) and shed light onto the possible mechanisms that might account for the experimental findings.

Jonikaitis et al. (2013) conducted two slightly different variants of their experiment. In one the attentional cue was turned off during saccade (task with transient cue), in the other the cue was shown until the end of the experiment (task with sustained cue). They determined the locus of attention by measuring performance in a discrimination task before and after the saccade at three different positions: the attention position (AP) where the attentional cue was presented; the lingering attention position (LAP), that is AP shifted by the saccade vector; and the remapped attention position (RAP), that is AP shifted by the reverse saccade vector. For both tasks, they found an attentional benefit at AP and RAP before saccade onset as well as an attentional benefit at AP and LAP after the saccade. They also observed a small but significant attentional benefit at RAP after saccade in the sustained cue task, which did not appear in the transient cue task. Since both transient and sustained cues led to similar attentional effects, it is possible that attention is largely driven by cue onset. Further, previous recordings in V4 show a strong decay in activation of a permanently present stimulus (Fischer & Boch, 1985). Thus, we simulated the study using a brief appearance of the attention cue. We used the same spatial setup as Jonikaitis et al. (2013): The model executes an 8° saccade to the right (modeled as described above), lasting 53 ms. Before saccade onset, attention is directed at the attention position (AP) located 6° above the fixation point due to a cuing stimulus at this position shown 180ms before saccade onset for 10ms. To account for the latency in the visual pathway, the activation of *Xr* in the model starts 50ms after stimulus onset. The spatial layout of the input signals, that are eye position, corollary discharge and retinal signal, are shown in Figure 4.

**Figure 4:**
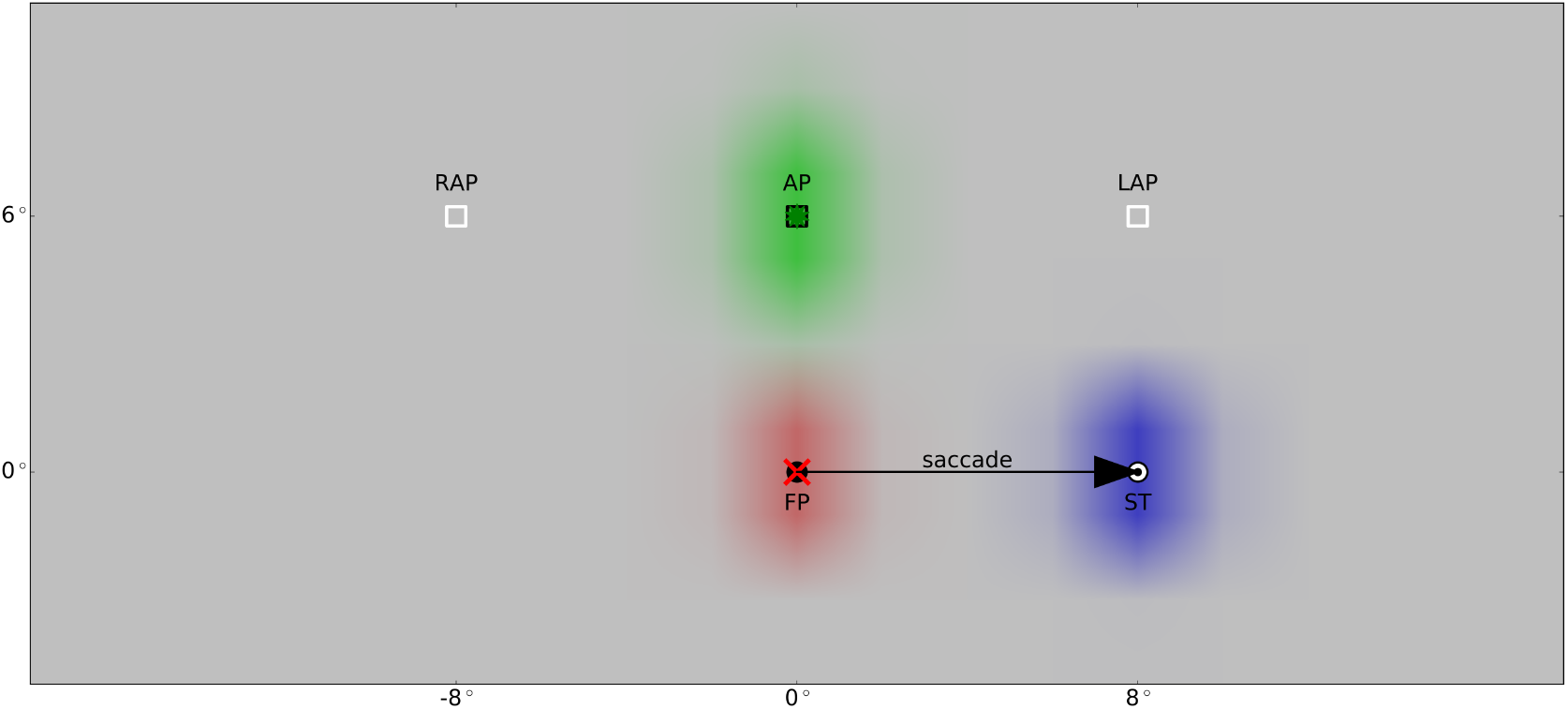
Setup for spatial updating of attention with cued attention with respect to the three input signals (cf. Figure 1): 180ms before saccade onset, a stimulus (green star) is presented at position AP at (0°, 6°) for 10ms to cue attention. The retinal signal (green) is delayed for 50ms to mimic the latency. After fixating the fixation point (FP) at (0°, 0°), the eyes reach the saccade target (ST) at (8°, 0°) in 53ms. The red cross symbolizes the current eye position. The eye position signal (red) starts to update 32ms after saccade offset to the new location of the eyes. The CD signal (blue) rises 86ms prior to saccade onset and is active for 171ms with its peak activity reached 10ms after saccade onset. Place markers for the remapped and the lingering attention position (RAP at (*−*8°, 6°) and LAP at (8°, 6°)) are shown. As the retinal signal and the origin of the corollary discharge signal is retinotopic, it shifts with the eye movement. In contrast, the PC signal is head-centered and therefore fixed during the saccade.

While Jonikaitis et al. (2013) used a behavioral paradigm to estimate the amount of attention distributed accross saccade, we directly plot the LIP activity as a measure of attention. Figure 5 shows the activity of both maps at different times. Again, since the LIP maps are four-dimensional, for visualization we projected them to two two-dimensional planes representing either the horizontal or the vertical information of this map. In our model, each LIP map triggers an attention pointer in a retinotopic reference frame. At the beginning, before the saccade, the only inputs to the model are eye position (PC) signal encoding the current eye position at FP and retinal signal to *Xr* encoding the cued attention position at AP. The retinal signal is combined multiplicatively with the PC signal in LIP PC which leads to a single activation blob (at (0°, 0°) for horizontal and (0°, 6°) for vertical, respectively). Projecting the activity of the LIP map back to visual space shows the attention pointer (red blob) at the desired attention position as shown in Figure 5A. Additionally, the cue generates an activity line in the second LIP map *Xb_CD_* which leads to an attention pointer at AP, too (covered by the red blob). Meanwhile, both LIP maps interact with each other via the neurons of *Xh*. All activity in *Xb_PC_* and *Xb_CD_* is summed up along each diagonal, fed into *Xh* and projected back along the same diagonal. Thus, the activity of *Xh* is projected back into *Xb_PC_* along the diagonal where it is combined multiplicatively with the PC signal. This sustains the same position as initially triggered by the cue. Importantly, as the saccade is being planned, the CD signal rises. It feeds into *Xb_CD_* and is multiplicatively combined with the reentrant signal from *Xh* feeding in along the diagonal. This leads to an activity blob in LIP CD (at (8°*, −*8°) for horizontal and (0°, 6°) for vertical, respectively). Projected to visual space, this activity triggers a second attention pointer at the remapped attention position (RAP) 8° to the left of the attention position (see Figure 5B, blue blob). This means that shortly before saccade onset, there are two spatial locations that exhibit attentional facilitation: The remapped attention position and the attention position itself. During saccade, the triggered attention pointers are shifted along with the eye movement as they are encoded in a retinotopic reference frame. Thus, after the saccade the attention pointers are shifted by 8° to the right. Therefore, the attention pointer induced by *Xb_PC_* is now at the lingering attention position (LAP) and the attention pointer induced by *Xb_CD_* is at AP (Figure 5C). As the CD signal decays after saccade onset, the activity in *Xb_CD_* also decays and the attention pointer induced by this map is gradually removed until it is completely extinguished. After the eyes landed at the saccade target (ST), the PC signal updates to the new eye position, thus the activity in *Xb_PC_* updates as well (to (8°*, −*8°) for horizontal and (0°, 6°) for vertical, respectively), and the attention pointer triggered by this map is remapped back to the attention position AP (see Figure 5D).

**Figure 5:**
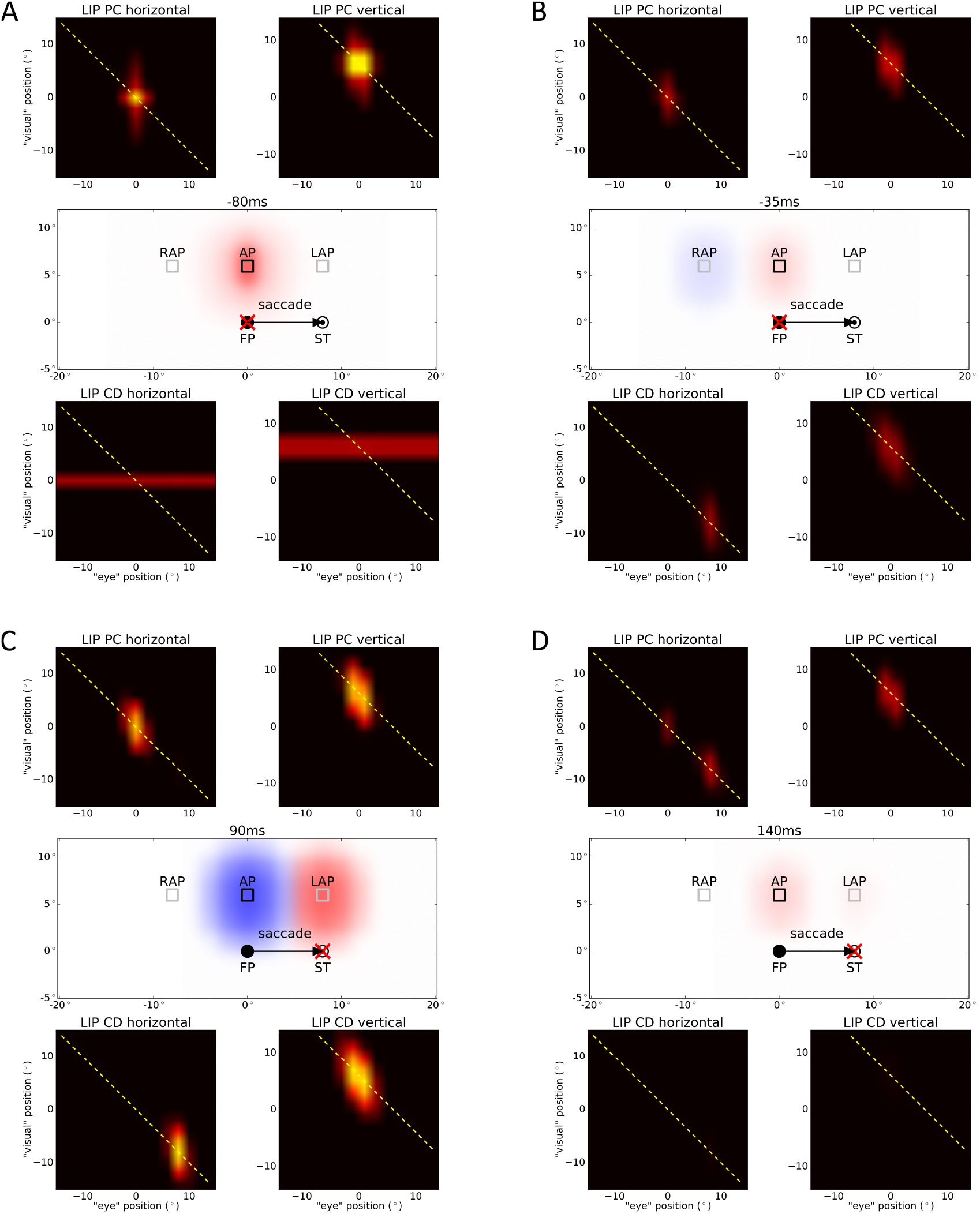
Simulation results of spatial updating of attention with cued attention for different time steps. The activities of both LIP maps (*Xb_PC_, Xb_CD_*) as well as the setup and the triggered attention pointers are plotted. Both LIP maps interact with each other by summing up all activations along each diagonal towards each neuron in *Xh*, and projecting back along the same diagonal. The yellow dashed line shows the diagonal which has the largest activation. The symbols are identical to Figure 4. The time in ms is aligned to saccade onset. (A) Long before saccade, the attention pointer at the desired attention position (AP) is encoded by the LIP map for the PC signal (red blob) and by the LIP map for the CD signal (blue blob, coved by red blob). (B) Shortly before saccade, the CD signal rises and activates neurons in LIP CD, which trigger a second attention pointer at the remapped attention position (RAP, blue blob). (C) Shortly after saccade, both attention pointers are shifted according to the eye movement as they are retinotopic. This leads to an attention pointer at the lingering attention position (LAP, red blob) and another one at AP (blue blob). (D) Long after saccade, the PC signal updates to the new eye position (ST) and the CD signal decays, so there is again only one attention pointer triggered by LIP PC at AP (red blob).

Figure 6 shows the time course of attention at AP (attention position), RAP (remapped attention position), and LAP (lingering attention position) over time. The three panels show the effect separated for the two LIP maps (red and blue lines, respectively) as well as the sum of the activities of both LIP maps (purple dashed lines) for each position. RAP is attended only before saccade onset generated by LIP CD cells. In contrast LAP is only attended after saccade offset through an attention pointer maintained by LIP PC cells. AP receives attention over the whole period, as before saccade, attention at AP is triggered by LIP PC and LIP CD and after saccade, the origin of the attention pointer switches from LIP CD to LIP PC.

**Figure 6:**
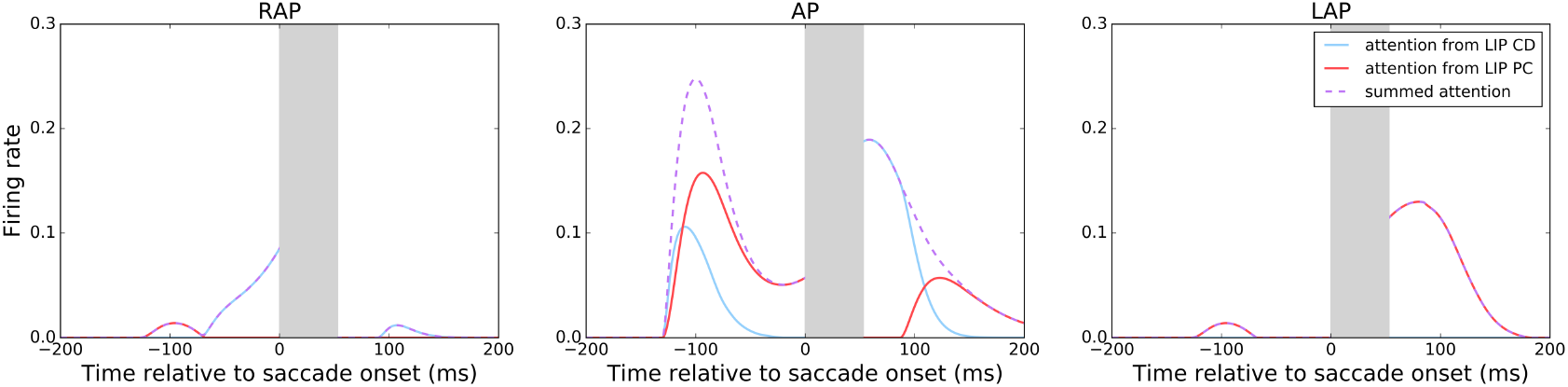
Attentional effect at different spatial positions over time for cued attention. The three plots show the attentional effect at the three spatial positions RAP (remapped attention position, left), AP (attention position, middle), and LAP (lingering attention position, right) over time. The red lines mark attention triggered by LIP PC, the blue lines mark attention triggered by LIP CD. The purple dashed lines mark the overall attention at this position. The gray bar covers the time of eye movement. The time in ms is aligned to saccade onset.

In the top-down mode we cue the desired location of attention to the model by an endogenous signal via *Xh*. This top-down attention signal may encode a head-centered reference position kept in memory. The emergence of this signal is not explicitly modeled here, but it might arise from memory recall in areas such as the Medial Temporal Lobe (Byrne, Becker, & Burgess, 2007). To demonstrate that this version of the model equally well explains the observations of Jonikaitis et al. (2013), we simulated the same experiment as before, but instead of a flashed cuing stimulus, we introduced top-down attention directed to the attention position as a constant input to *Xh* (see Figure 1). The spatial layout of the input signals, here eye position, corollary discharge and top-down attention, are shown in Figure 7.

**Figure 7:**
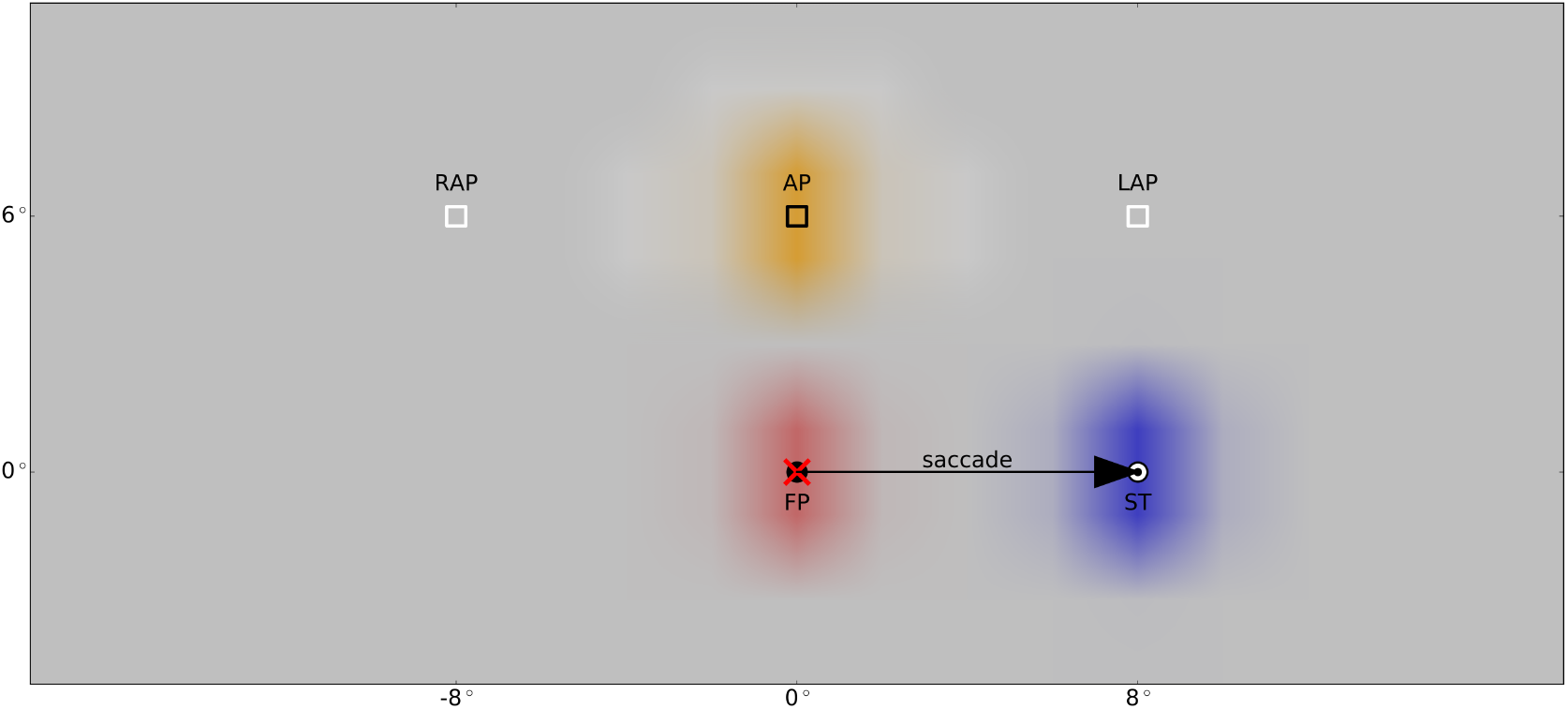
Setup for spatial updating of attention with top-down attention with respect to the three input signals (cf. Figure 1): After fixating the fixation point (FP) at (0°, 0°), the eyes reach the saccade target (ST) at (8°, 0°) after 53ms. The red cross symbolizes the current eye position. The eye position signal (red) starts to update 32ms after saccade offset to the new location of the eyes. The CD signal (blue) rises 86ms prior to saccade onset and is active for 171ms with its peak activity reached 10ms after saccade onset. During the whole process, a top-down attention signal (orange) is introduced at position AP at (0°, 6°). The corollary discharge signal is retinotopic, thus, it shifts with the eye movement. In contrast, PC signal and attention signal are head-centered and therefore fixed during the saccade. Markers indicate the remapped and the lingering attention position (RAP at (−8°, 6°) and LAP at (8°, 6°)).

Figure 8 shows the activity of both maps at different times. At the beginning, before the saccade, the only inputs to the model are eye position (PC) signal encoding the current eye position at FP and top-down attention signal to *Xh* encoding the attention position at AP. The attention signal is projected backwards into the LIP map *Xb_PC_* along the diagonal and is combined multiplicatively with the PC signal which leads to a single activation blob (at (0°, 0°) for horizontal and (0°, 6°) for vertical, respectively). This activity maintains an attention pointer (red blob) at the desired attention position as shown in Figure 8A. Shortly before saccade onset, the CD signal rises and feeds into the second LIP map, *Xb_CD_*, where it is multiplicatively combined with the attention signal from *Xh* feeding in along the diagonal. This leads to an activity blob in LIP CD (at (8°*, −*8°) for horizontal and (0°, 6°) for vertical, respectively) and a second attention pointer at the remapped attention position (RAP) 8° to the left of the attention position (see Figure 8B, blue blob). Thus, like in the previous simulation, the two attention pointers triggered by the LIP maps direct attention to the remapped attention position and the attention position itself shortly before saccade onset. Likewise, shortly after saccade the attention pointers are shifted to AP and LAP, respectively (see Figure 8C) and long after the saccade, when the CD signal has decayed and the PC signal is updated to the saccade target, there is again only one attention pointer triggered by LIP PC directing attention to AP (see Figure 8D).

**Figure 8:**
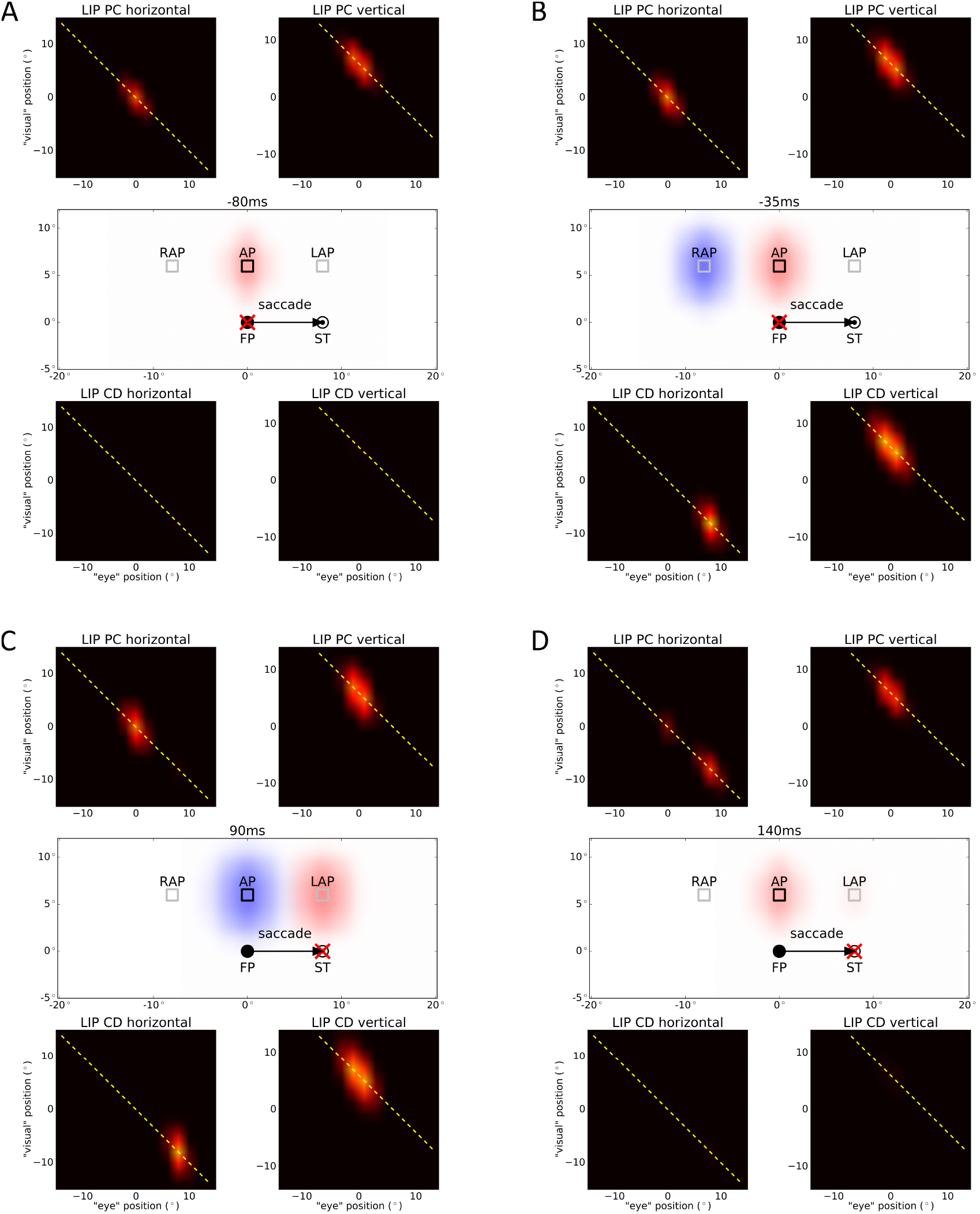
Simulation results of spatial updating of attention with top-down attention for different time steps. The activities of both LIP maps (*Xb_PC_, Xb_CD_*) as well as the setup and the triggered attention pointers are plotted. Additionally, the diagonal on which the attention signal is fed into the LIP maps is plotted with a yellow dashed line. The symbols are identical to Figure 7. The time in ms is aligned to saccade onset. (A) Long before saccade, the attention pointer at the desired attention position (AP) is encoded by the LIP map for the PC signal (red blob). (B) Shortly before saccade, the CD signal rises and activates neurons in LIP CD, which trigger a second attention pointer at the remapped attention position (RAP, blue blob). (C) Shortly after saccade, both attention pointers are shifted according to the eye movement as they are retinotopic. This leads to an attention pointer at the lingering attention position (LAP, red blob) and another one at AP (blue blob). (D) Long after saccade, the PC signal updates to the new eye position (ST) and the CD signal decays, so there is again only one attention pointer triggered by LIP PC at AP (red blob).

## 4 Discussion

Our simulation results suggest that predictive remapping of receptive fields and spatial updating of attention are two sides of the same coin. Predictive remapping in our model arises from a relatively small, and simple, recurrent neural circuit. Although the model likely simplifies from the biophysical implementation in the brain, much can be explained at this abstract level. The remapping results from the interaction of the phasic corollary discharge signal and the tonic feedback from LIP, which means it is synchronized with the CD signal, i.e. saccade onset. The CD signal has been recently well explored in electrophysiological studies (Sommer & Wurtz, 2004, 2008). The model suggests that this signal is integrated into LIP gain fields. As the CD signal contains information about the future position of the eyes, it contributes to a new reference to which the flashed stimulus is anchored. Thus, predictive remapping of receptive fields can be well understood by the integration of a second spatial reference system within a gain-field like representation. As the gain-fields operate in a retinocentric coordinate frame and the internal eye position updates after saccade, the flashed stimulus is transferred against saccade direction due to the future eye position reference, leading to the activation of LIP neurons at the RF location if the stimulus has been flashed at FRF. From the viewpoint of these LIP neurons, it looks like their RFs have shifted forward.

Regardless of the particular experimental paradigm, around saccades our model suggests the presence of two pointers linked to different eye related signals, one to a proprioceptive eye position and the other to the corollary discharge. In the spatial attention task, the corollary discharge signal leads to a retinotopic attention pointer at the remapped attention position shortly before saccade onset, which is shifted with the saccade to the original position of attention and is finally removed shortly after the eyes have landed at the saccade target consistent with Rolfs et al. (2011). Furthermore, the model explains the lingering of attention (Golomb et al., 2008, 2010) by a late-updating, proprioceptive eye position signal which triggers a retinotopic attention pointer first at the desired position. This pointer is then shifted by the saccade and finally, with the updated signal, remapped to the initial position without covering intermediate positions (Golomb, Marino, Chun, & Mazer, 2011). Hence, we have two attention pointers generated by LIP that are retinotopic. These pointers are shifted during the eye movement according to the saccade vector. Thus, during saccades there is one pointer which moves from the remapped attention position towards the actual attention position and one pointer moving away from the attention position. Consequently, while the eyes are moving, there is no attention directed to the original attention position itself as shown by Yao, Ketkar, Treue, and Krishna (2016). Furthermore, Yao et al. (2016) conclude from their experimental results that the attention position must be attended again within 30ms after the eyes have landed. In our model, attention is available at the attention position immediately after the saccade as the remapped attention pointer is then shifted to this position. Taking into account that the decision process in the experiment itself needs time (processing the visual input and making the decision), this fits to the data of Yao et al. (2016). In addition to attentional cuing, we showed that attention may also be induced by an endogenous attention signal.

The functional roles of the two eye-related signals used in our model have been recently discussed in a review (Sun & Goldberg, 2016). As the (proprioceptive) eye position signal is inaccurate after saccade, LIP gain fields may not be suitable to solve the spatial accuracy problem. Therefore, Sun and Goldberg (2016) conclude, that there are “[…] two different representations of space: a rapid retinotopic one and a slower craniotopic one.”. In the model presented here, these two representations interact to trigger predictive remapping and spatial updating of attention. Both phenomena can be well explained by a lateral or reentrant network at the level of LIP, such that the presaccadic activation is re-computed in the future reference frame by means of the corollary discharge. For simplicity, we assumed two different neuron types, CD and PC eye-related cells. However, it is likely that the separation is not so complete and there may be a continuum of cells.

Since previous simulations with the 1D model version indicate that mislocalization in total darkness around saccade and saccadic suppression of displacement also can be accounted for by the same neural circuits (Ziesche & Hamker, 2011, 2014; Ziesche et al., 2017) and as the 1D version of this model was designed prior to the observation of updating of attention, from the models point of view this observation can be considered an inherent prediction.

Although the newly presented and the previous 1D model does a good job of accounting for several outstanding issues and accurately replicates the behavioral findings of Jonikaitis et al. (2013), there are nevertheless a number of issues that have to be addressed in future work. First, the exact timing of the proprioceptive eye position signal needs to be explored in more detail. While a recent study in LIP suggests that gain fields may update even after 150ms (B. Y. Xu, Karachi, & Goldberg, 2012), which is about 80 − 100ms later than in our model, Y. Xu et al. (2011) report an update after 60ms in somatosensory cortex. No such information exists in humans. Our parameters are based on previous versions of the model (Ziesche & Hamker, 2011, 2014; Ziesche et al., 2017) and mainly motivated to account for human behavioral data. Second, there is some variation in experimental studies regarding the dominance of the effects in the data which requires future clarifications. For instance, Marino and Mazer (2018) find in V4 only evidence for predictive remapping of attention but not for lingering of attention whereas Yao, Treue, and Krishna (2018) detect in MT only lingering but no remapping of attention. Yao et al. (2016) even find neither remapping nor lingering of attention in a psychophysical study with humans though these results are may be biased by their experimental design where remapped and lingering attention position were not only irrelevant, but importantly these positions were never target locations that tested the amount of attention. Lisi, Cavanagh, and Zorzi (2015) suggest, that spatial updating of attention depends on the setup of the experiment, namely whether visual objects, which can serve as spatial landmarks, are presented or not. In their experiments, they found that the lingering effect vanishes if a placeholder is shown at the attended location for the whole trial. However, the ability to maintain attention at a spatiotopic location during saccades increases with the presence of placeholders. These findings may help to classify the various contradictory results.

## Acknowledgments

This work has been supported by the US-German collaboration on computational neuroscience (BMBF 01GQ1409) and partly supported by the European Union’s Seventh Framework Programme (FET, Neuro-Bio-Inspired Systems: Spatial Cognition) under grant agreement no. 600785. We thank J. Mazer for his comments and discussions.

## Appendix Neuro-computational Model

The neurons in each map follow different ordinary differential equations (ODEs). In our extension of the original 1D model to the two-dimensional one, we use the same ODEs as stated in Ziesche and Hamker (2011), but with some simplifications and extensions. For all ODEs that compute firing rates *r*, we set negative values to zero.

- Firing rates of neurons in map *Xr* representing the stimulus position in a eye-centered reference frame:

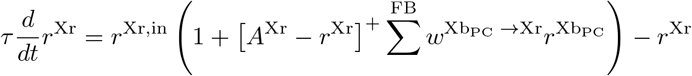

with *r*^Xr,in^ the sensory bottom-up input created by a given stimulus:

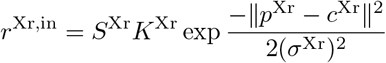 Here, *S*^Xr^ is the short-term synaptic depression simulated as in Hamker (2005) modeling the decaying response strength over time while the stimulus is presented, *K*^Xr^ defines the strength of the stimulus and 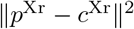 is the distance between stimulus position *p*^Xr^ and receptive field center *c*^Xr^ for each neuron of the map. As we now have a two-dimensional visual scene, the stimulus position as well as the receptive field center are two-dimensional.
- Firing rates of neurons in map *Xe_PC_* and *Xe_CD_* representing eye position in head and retinotopic eye displacement, respectively:

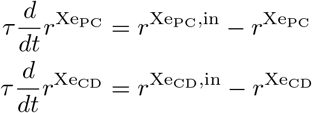

where 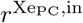 and 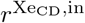, in are Gaussian input signals modeling the proprioceptive eye position signal and the corollary discharge signal, respectively:

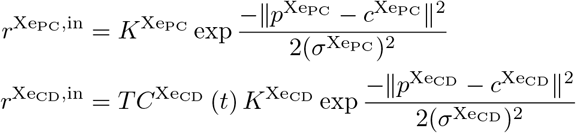 *K^Xe_PC_^* and *K^Xe_CD_^* are the strengths of the corresponding signal, 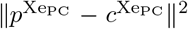 is the distance between eye position *p^Xe_PC_^* and center of eye position tuning *c^Xe_PC_^* for each neuron of map *Xe_PC_* and likewise, 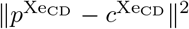 is the distance between eye displacement *p^Xe_CD_^* and center of eye displacement tuning *c^Xe_CD_^* for each neuron of map *Xe_CD_*. Again, the positions are now two-dimensional. *TC^Xe_CD_^(t)* models the time course of the phasic corollary discharge signal, namely rise and decay around saccade onset:

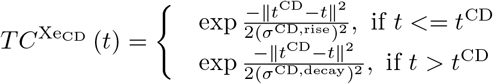

with *t*^CD^ the time, where the CD signal reaches its maximum. In our model, this maximum is at 10ms after saccade onset consistent with data of Ferraina, Paré and Wurtz (2002).
- Firing rates of neurons in map *Xe_FEF_* representing the eye displacement in a head-centered reference frame:

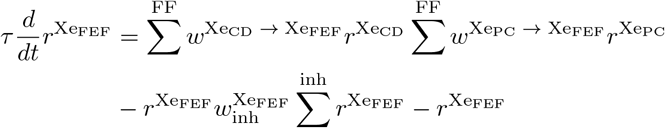 In contrast to the original ODE, we simplified the firing rate for *Xe_FEF_* by removing the saturation term as well as the gain modulation term. Thus, this map is now a classical basis function map instead of a gain modulation map to combine PC and CD signal.
- Firing rates of neurons in map *Xb_PC_* representing the joint representation of stimulus position and eye position:

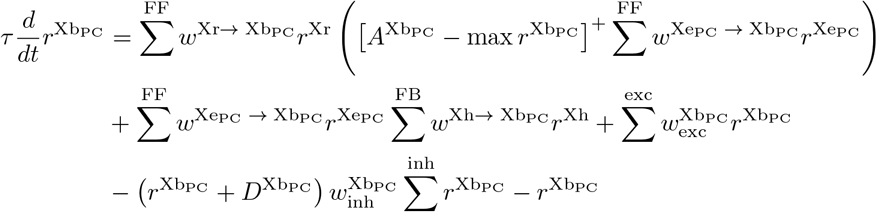 For the firing rates of *Xb_PC_* we added an additional feedback signal that combines the PC signal with the signal of the intermediate cells of *Xh*. This feedback signal is identical to the feedback signal of the other LIP map *Xb_CD_*. Additionally, we removed the perisaccadic suppression factor on the input from *Xe_PC_* as otherwise the feedback from *Xh* would be reduced over a certain period of time due to the multiplicative interaction with the PC signal.
- Firing rates of neurons in map *Xb_CD_* representing the joint representation of stimulus position and eye displacement:

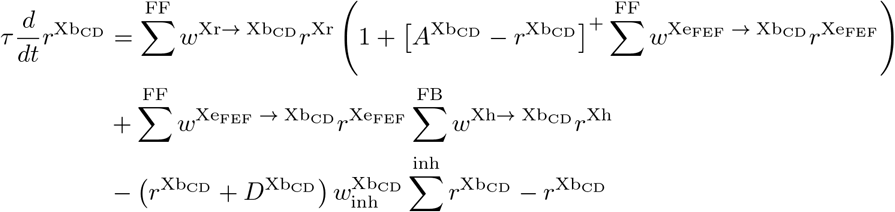
- Firing rates of neurons in map *Xh*:

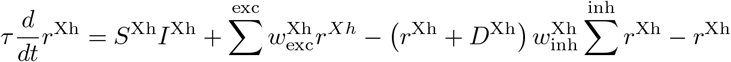

with *I*^Xh^ the input consisting of the feedforward input from both LIP maps and a newly introduced, attentional top-down signal *r*^Xh,in^:

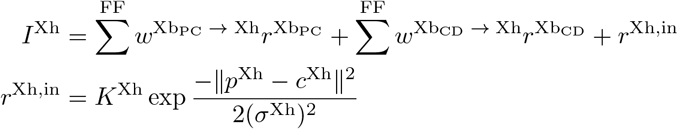

where *K*^Xh^ denotes the strength of the attention and ||*p*^Xh^ −*c*^Xh^||^2^ the distance between attention position *p*^Xh^ and center of attention position tuning *c*^Xh^ for each neuron of map *Xh*, each with two dimensions. *S*^Xh^ is the synaptic suppression simulated as in the study by Hamker (2005):

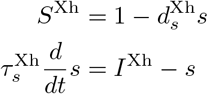

As the dimension of each map has to be doubled compared to the definition of the maps in Ziesche and Hamker (2011), we reduced the number of neurons in each dimension to compensate for the higher computational effort. More precisely, we have 21 neurons for the horizontal and 15 neurons for the vertical covering a rectangular visual field of 40° *×* 30°. Thus, *Xr, Xe_PC_, Xe_CD_* and *Xh* are now two-dimensional containing 21 *×* 15 neurons.

The connections between the different maps are defined through the different weights used in the ordinary differential equations. The connection weights follow Gaussian functions dependent on the distance between the position of the neuron in the pre-synaptic map and the position of the neuron in the post-synaptic map, i.e.
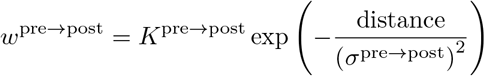

with *w*^pre*→*post^ set to 0 if the value is lower than 0.001. The measurement of the distance differs among the connections due to the different dimensions of the maps and the different ways of connecting maps. There are three types of connections: horizontal, vertical and diagonal. The horizontal and vertical connections are used to connect two-dimensional maps with four-dimensional maps where two of the four dimensions are disregarded (either the first two or the last two). For example, we connect *Xr* with *Xb_PC_* horizontally independent of the last two dimensions of *Xb_PC_*. That means, the distance used for the Gaussian function depends only on the position of the neuron in *Xr* and the first two position parameters of the neuron in *Xb_PC_*. More precisely, suppose we want to connect neuron (*i, j*) of *Xr* with neuron (*k, l, m, n*) of *Xb_PC_*. The distance between these neurons is then calculated by:

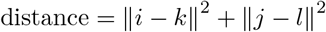

The weight between neuron (*i, j*) and neuron (*k, l, m, n*) is then:

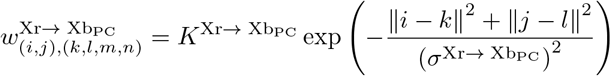

This connection pattern is also used to connect *Xr* with *Xb_CD_*, *Xe_CD_* with *Xe_FEF_* and *Xb_PC_* with *Xr*.

Similarly, a vertical connection between a two-dimensional map and a four-dimensional map makes only use of the last two position parameters of the neurons in the four-dimensional map to calculate the distance. Thus, the distance between neuron (*i, j*) of a two-dimensional map and neuron (*k, l, m, n*) of a four-dimensional map is then:

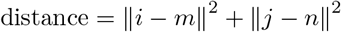

Such a vertical connection is used to connect *Xe_PC_* with *Xe_FEF_* and to connect *Xe_PC_* with *Xb_PC_*.

These definitions of the connection patterns allows the interpretation of the four dimensions of a map as follows: The first two dimensions represent the horizontal and the vertical information of a horizontally connected input and the last two dimensions represent the horizontal and the vertical information of a vertically connected input.

For a diagonal connection pattern we use both horizontal and vertical information of the four-dimensional map to connect it with a two-dimensional map. We use such a connection pattern to connect *Xh* with the LIP maps *Xb_PC_* and *Xb_CD_* and vice versa. The distance between neuron (*i, j*) of *Xh* and neuron (*k, l, m, n*) of a LIP map is defined as follows:

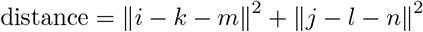

For the remaining connection pattern between *Xe_FEF_* and *Xb_CD_* that connects two four-dimensional maps, we read out the pre-synaptic map diagonally and connect this vertically with the post-synaptic map, i.e. we use all four dimensions of *Xe_FEF_* and only the last two dimensions of *Xb_CD_* to calculate the distance between the neurons. More precisely, if we want to connect neuron (*i, j, k, l*) of *Xe_FEF_* with neuron (*m, n, o, p*) of *Xb_CD_*, the distance between these neurons is:

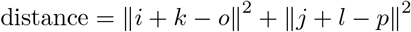

The lateral excitatory connections used in map *Xb_PC_* and *Xh* are defined as Gaussian functions dependent on the distance between the positions of the neurons in the maps considering all dimensions of the map. Thus, the calculation of the weights follows the equation:

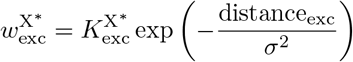

with

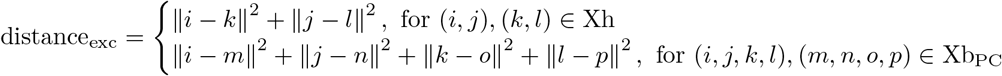

To reduce the computational effort, the lateral inhibitory connections with fixed weights 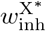 are not created as explicit connections, but calculated with the help of the mean over all firing rates multiplied by the total number of neurons:

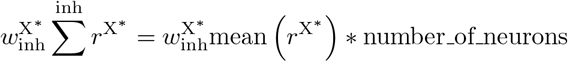

At last, Table 1 lists all parameters whose values have changed in comparison to Ziesche and Hamker (2011). Mainly, the adaption of the values are due to the three major changes in the model: reduction of number of neurons, simplifying *Xe_FEF_*, and adding feedback from *Xh* to *Xb_PC_*. The new parameter values for this feedback connection are listed in the last two rows and are equal to those for the feedback connection from *Xh* to *Xb_CD_*.

**Table 1:** Parameters whose values have changed in comparison to Ziesche and Hamker (2011). The first column contains the name of the parameter, the second column the equation number in Ziesche and Hamker (2011) dealing with this parameter. The third and fourth column state the original and the new value of the parameter. The last two rows contain new parameters.

| parameter | equation | original value | new value |
| --- | --- | --- | --- |
| $\sigma^{PC}$ | 5 | 8.0° | 1.0° |
| $\sigma^{Xr \rightarrow Xb_{PC}}$ | 9 | 1.0° | 0.5° |
| $K^{Xr \rightarrow Xb_{PC}}$ | 9 | 0.6 | 2.0 |
| $\sigma^{Xb_{PC} \rightarrow Xr}$ | 10 | 1.0° | 0.5° |
| $\sigma^{Xe_{PC} \rightarrow Xb_{PC}}$ | 11 | 10.0° | 2.0° |
| $K^{Xe_{PC} \rightarrow Xb_{PC}}$ | 11 | 10.0 | 1.0 |
| $w_{inh}^{Xb_{PC}}$ | 12 | 0.4 | 0.04 |
| $\sigma_{exc}^{Xb_{PC}}$ | 13 | 1.0° | 0.5° |
| $\sigma^{CD}$ | 14 | 8.0° | 1.0° |
| $\sigma^{CD, rise}$ | 16 | 50.0 | 65.0 |
| $\sigma^{CD, decay}$ | 17 | 150.0 | 50.0 |
| $\sigma^{Xe_{CD} \rightarrow Xe_{FEF}}$ | 18 | 1.0° | 0.5° |
| $K^{Xe_{CD} \rightarrow Xe_{FEF}}$ | 18 | 0.5 | 5.0 |
| $\sigma^{Xe_{PC} \rightarrow Xe_{FEF}}$ | 19 | 2.0° | 1.0° |
| $K^{Xe_{PC} \rightarrow Xe_{FEF}}$ | 19 | 15.0 | 10.0 |
| $\sigma^{Xr \rightarrow Xb_{CD}}$ | 21 | 1.0° | 0.5° |
| $\sigma^{Xe_{FEF} \rightarrow Xb_{CD}}$ | 22 | 1.0° | 0.5° |
| $K^{Xe_{FEF} \rightarrow Xb_{CD}}$ | 22 | 3.0 | 20.0 |
| $\sigma^{Xh \rightarrow Xb_{CD}}$ | 23 | 45.0° | 2.0° |
| $K^{Xh \rightarrow Xb_{CD}}$ | 23 | 0.13 | 1.3 |
| $w_{inh}^{Xb_{CD}}$ | 25 | 0.2 | 0.02 |
| $\sigma^{Xb_{PC} \rightarrow Xh}$ | 28 | 15.0° | 7.5° |
| $K^{Xb_{PC} \rightarrow Xh}$ | 28 | 0.35 | 0.015 |
| $\sigma^{Xb_{CD} \rightarrow Xh}$ | 29 | 15.0° | 7.5° |
| $K^{Xb_{CD} \rightarrow Xh}$ | 29 | 0.2 | 0.015 |
| $w_{inh}^{Xh}$ | 30 | 1.0 | 0.1 |
| $\sigma_{exc}^{Xh}$ | 30, 34 | 1.0° | 0.5° |
| $w_{exc}^{Xh}$ | 30, 34 | 0.2 | 0.4 |
| $\sigma^{Xh \rightarrow Xb_{PC}}$ | — | — | 2.0° |
| $K^{Xh \rightarrow Xb_{PC}}$ | — | — | 1.3 |

